# Characterization of cell-cell communication in COVID-19 patients

**DOI:** 10.1101/2020.12.30.424641

**Authors:** Yingxin Lin, Lipin Loo, Andy Tran, Cesar Moreno, Daniel Hesselson, Greg Neely, Jean Y.H. Yang

## Abstract

COVID-19 patients display a wide range of disease severity, ranging from asymptomatic to critical symptoms with high mortality risk. Our ability to understand the interaction of SARS-CoV-2 infected cells within the lung, and of protective or dysfunctional immune responses to the virus, is critical to effectively treat these patients. Currently, our understanding of cell-cell interactions across different disease states, and how such interactions may drive pathogenic outcomes, is incomplete. Here, we developed a generalizable workflow for identifying cells that are differentially interacting across COVID-19 patients with distinct disease outcomes and use it to examine five public single-cell RNA-seq datasets with a total of 85 individual samples. By characterizing the cell-cell interaction patterns across epithelial and immune cells in lung tissues for patients with varying disease severity, we illustrate diverse communication patterns across individuals, and discover heterogeneous communication patterns among moderate and severe patients. We further illustrate patterns derived from cell-cell interactions are potential signatures for discriminating between moderate and severe patients.

## Introduction

Infection with SARS-CoV-2 causes COVID-19 and is driving the ongoing pandemic impacting the global population. The respiratory system is the primary route of infection for SARS-CoV-2, due to a combination of airborne transmission (*1*) and the presence of the SARS-CoV-2 receptor ACE2 in human airways (*2*–*4*). SARS-CoV-2 infection causes a spectrum of symptoms. Patients can be asymptomatic, exhibit mild symptoms, or develop severe disease with increased risk of death (*5, 6*). Disease outcome is dictated by a combination of direct viral effects on patient tissues (*7*), protective antiviral immunity (*8*) and overexuberant antiviral or inflammatory immune responses driving tissue damage (*9, 10*). However, it is not clear why some patients experience mild symptoms while others succumb to the illness, and how communication between host compartments controls the disease progression.

On inoculation, SARS-CoV-2 can infect cells in the oral/nasal mucosa (*11*) and nasopharynx expressing low levels of ACE2 (*12*), with the virus descending to the lower airways in some patients to infect ACE2+ type II alveolar epithelial cells (*13*). In more severe disease, SARS-CoV-2 infected cells likely enhance or alter their communication networks to recruit additional immune support, in part through excessive local innate immune engagement (*14*) which can then recruit neutrophil and T cells from the blood which further escalate the inflammatory cascades leading to a “cytokine storm” in patients with severe disease (*15*), (*16*). This vicious cycle eventually drives pathological inflammatory cell and fluid accumulation and extensive tissue damage, leading to lung stiffness, impaired O_2_/CO_2_ exchange (together called acute respiratory distress syndrome; ARDS), and death. Paradoxically, many patients do exhibit lower airway infection/inflammation yet experience only mild disease, and it is unclear why these patients avoid critical disease.

To define the cellular transcriptional responses involved in COVID-19 severity, single-cell RNA-seq has been performed on patient samples, including peripheral blood mononuclear cells (PBMCs), bronchoalveolar lavage. These studies further reinforce the notion that excessive inflammation correlates with negative disease outcome (*17, 18*). Beyond cell identification (*19*), single-cell analysis can also be used to infer cell-cell interactions (*20*–*22*) and these approaches can help inform disease mechanisms.

In this study, we harness collections of single-cell COVID-19 data sets available to evaluate the molecular patterns associated with disease severity. Recognizing the importance of cell-cell communication networks within the infected lung, we develop a generalizable workflow to explore cell-cell interactions. We then apply our novel workflow to analyze the cell-cell communication networks from patients with varying disease severity and pinpoint critical cell types and cell-cell communication channels that mark healthy network communication as well as discriminate disease severity.

## Results

### Generalizable workflow to identify and measure cell-cell communication in individuals

We develop a generalizable workflow based on statistical learning strategies that allow us to visualize, identify and characterize cell-cell interaction patterns (Fig. 1A). The workflow begins with joint classification using scClassify (*19*) based on four reference datasets (see Material and Methods) to refine cell type annotations. Next, to partition cell heterogeneity, unsupervised clustering is performed on each annotated cell type to further define subgroups of cells with the potential to identify cellular subtypes associated with different disease progression. Cluster merging (*23*) is used here to prevent overclustering. Finally, we calculate a cell-cell interaction score/measure for each individual COVID-19 sample between different cellular subtypes. Applying this workflow to single-cell data with multiple individuals will generate a large matrix for each individual sample with columns representing cell types and rows representing ligand-receptor pairs (Fig. 1A). Each ligand-receptor pair is further grouped into different pathways to facilitate interpretation. Details of this workflow are described in the Materials and Methods section.

**Fig. 1:**
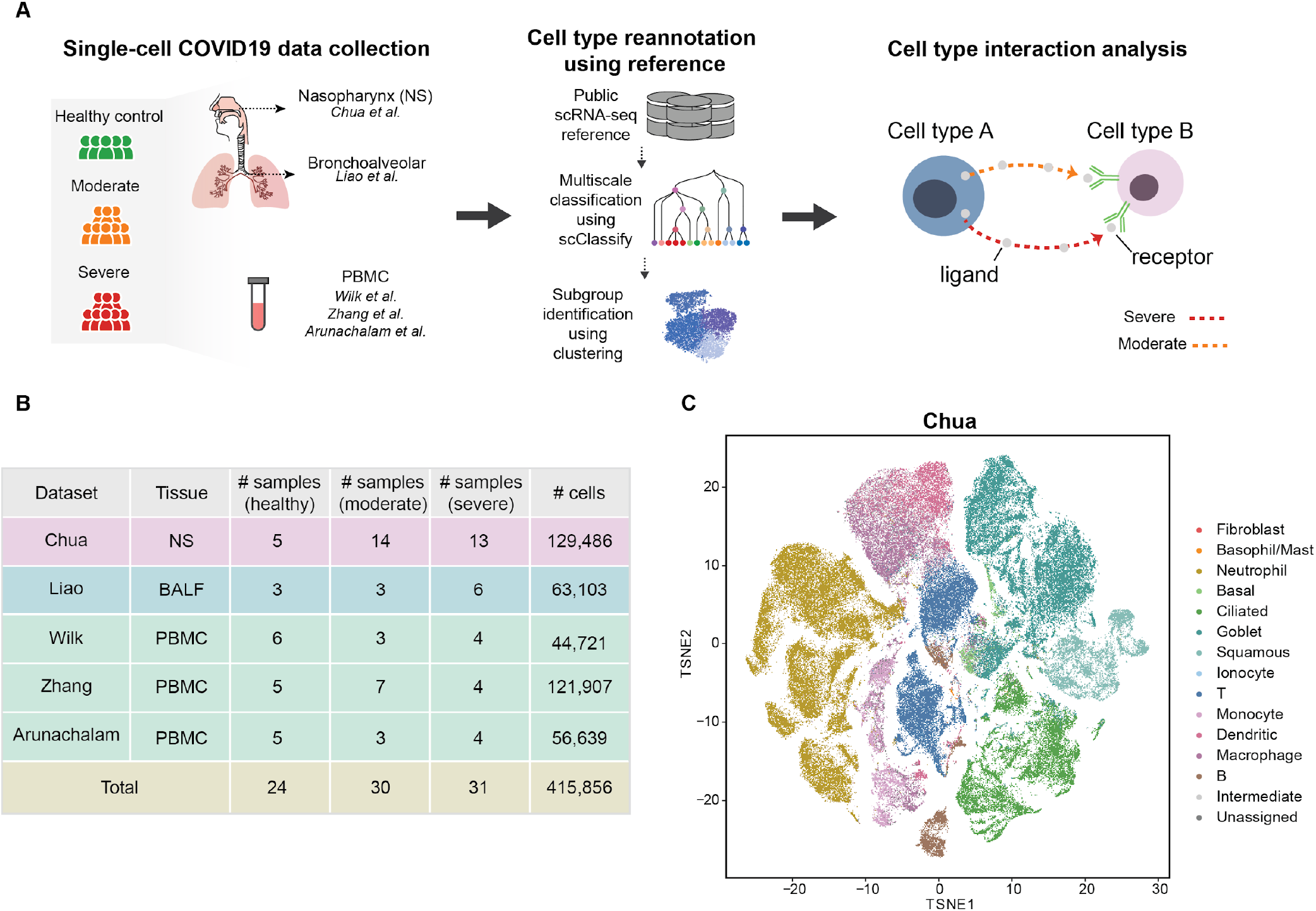
A.Schematic of the data analytic workflow. B. Summary of curated single-cell RNA-seq from COVID-19 studies from different tissues that are publicly available. C.tSNE plot illustrating cell types from all samples in the Chua dataset based on the reannotation using a modified version of the joint classification from scClassify built from four large reference datasets of human lungs.

We examined five publicly available single-cell RNA-seq datasets from COVID-19 patients with different degrees of severity, using samples from nasopharyngeal (NS), bronchoalveolar lavage fluid (BALF), and PBMCs, including a total of 415,856 cells from 24 healthy controls, 30 moderate and 31 severe samples (Fig. 1B). In NS and BALF tissues (*24, 25*), we re-annotated the cells using four healthy human lung scRNA-seq datasets (*4, 26, 27*) including 189,967 cells and 44 cell types. For the Chua dataset with 5 healthy controls, 14 moderate and 13 severe samples (Supplementary Fig. 1A-B), scClassify is able to identify additional cell types and provide further refinement as illustrated in Fig. 1C. For example, the original “outliers epithelial” cluster was refined to “ciliated cells”, and “secretory” cells to “goblet” cells. By accounting for such refinement, the new annotation recapitulates 78% of the original published analysis. The classified cell types are clearly identified by known markers (Supplementary Fig. 1C), and further clustering of the Chua dataset generates 50 subclusters. Similar reannotation is applied to the Liao dataset (Supplementary Fig. 2A-B) resulting in 15 cell types and 52 subclusters.

### Cell-cell interactions are significantly different in patients with COVID-19 compared to healthy individuals

To identify cell-cell interaction (CCI) differences with respect to disease severity in each dataset, we calculate the CCI scores that represent the communication probabilities among all pairs of subclusters across all ligand-receptor pairs (see Materials and Methods for details). Our group-specific CCI scores (CCI_group_) aggregate the scores across all different pathways between each major cell type pair for different disease severity groups, represented as a network graph with thicker edges indicating stronger cell-cell interaction. Our results highlight that in healthy controls, most cell-cell interactions are between basal, ciliated, and goblet cells of the lung epithelium, with dendritic cells providing immune surveillance (Fig. 2A). As disease severity increases, cell-cell interactions become dominated by interactions between the lung epithelium and proinflammatory players within the immune compartment (Fig. 2A-C, Supplementary Fig. 3A-E). Overall, we observe significantly less communication (fewer edges in Fig. 2A) in healthy individuals compared to moderate (Fig. 2B) and severe patients (Fig. 2C).

**Fig. 2:**
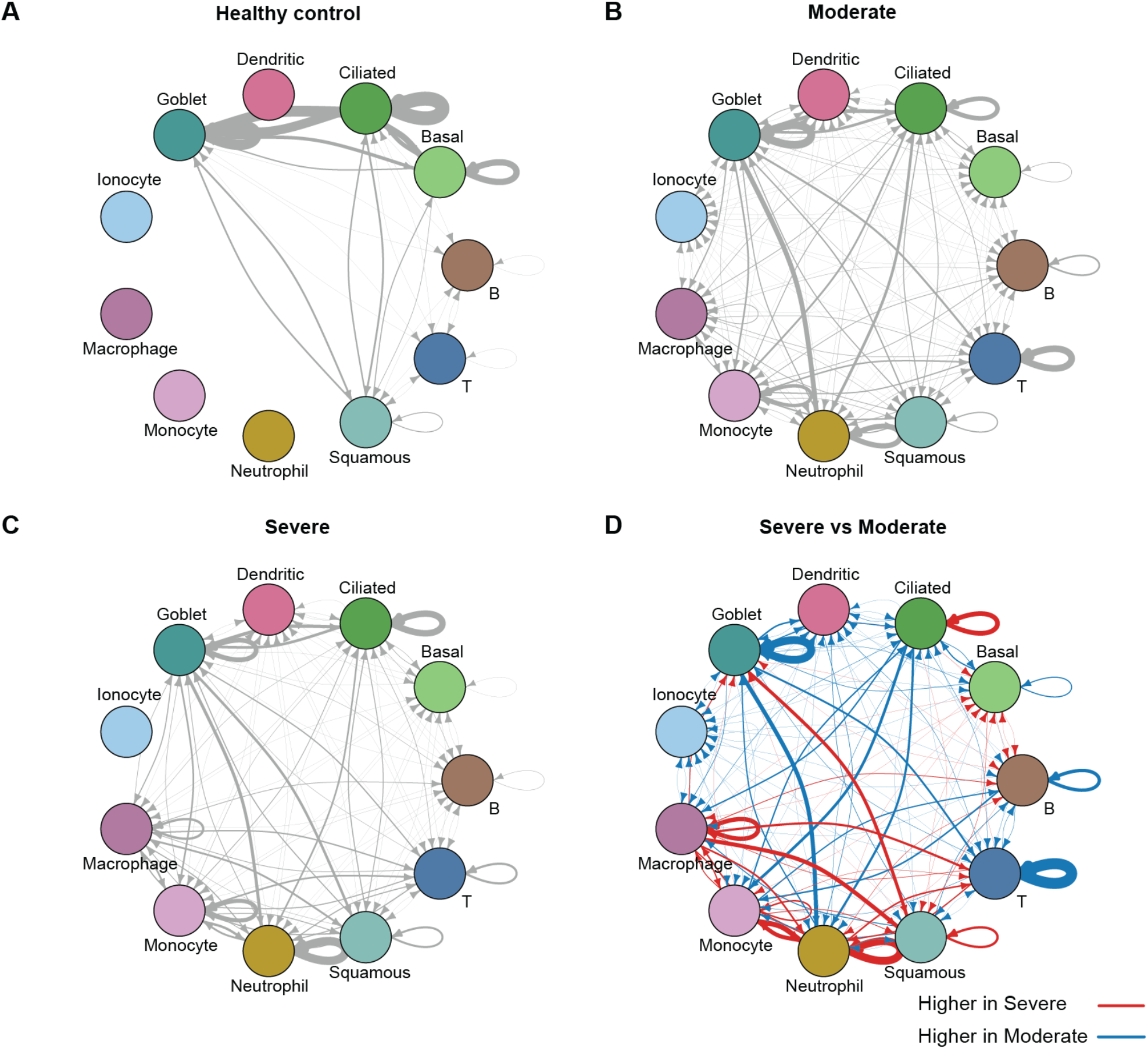
A - C. Network representing the group specific cell-cell interaction (CCI_group_) considering different disease severity as groups in the Chua dataset from (A) healthy controls (B) moderate patients and (C) severe patients. The nodes represent major cell types and the edges represent aggregate tCCI interaction signals across individuals from the same group. Thicker edges indicate stronger cell cell interaction signals. D. Network representing the difference of cell-cell interaction between severe and moderate patients. The nodes represent cell types and an edge measures the difference in cell-cell interaction. A red edge indicates an interaction higher in severe patients and a blue edge indicates an interaction higher in moderate patients.

We next study the cell-cell interaction in all three datasets (*18, 28, 29*). Applying our developed workflow, we unified the annotation across all three datasets (Supplementary Fig. 4A). We used the Wilk dataset (with 44,721 cells and 20 cell types) as a reference (*28*) to reannotate the cell types for the Zhang dataset and the Arunachalam dataset. Cells with the same annotation were well integrated with scMerge (*30*) (Supplementary Fig. 4A), indicating that similar cell types exist in the three studies. We observe distinct cell type compositions of these three studies, potentially due to different sampling or cell isolation procedures (Supplementary Fig. 4B). In all three PBMC studies, we observe that in monocyte-related interactions, COVID-19 patients generally have more cell-cell interactions than healthy controls (Supplementary Fig. 5A-C). The results across all five datasets suggest that cell type composition alone from single-cell experiments may not sufficiently discriminate between patients with different disease severity. This, along with the varying cell-cell interaction patterns across disease groups supports the further examination of the association between cell-cell interaction with disease outcomes and progression.

### Reduced T cell interactions and increased monocyte interactions in severe COVID-19 patients across different compartments

Focusing on immune cell interactions in the upper airway, we observe different cell type interaction patterns across patients with different disease severity (Fig. 2D, Supplementary Fig. 3F) in the Chua dataset. Compared to moderate patients, severe patients have less T cell to T cell interaction (illustrated by the thick blue edge on the T cell node in Fig. 2D) and more macrophage to macrophage interaction (marked by thick red edge on the macrophage node in Fig. 2D). Between different cell types, we see higher interactions between monocyte/macrophage or T cells towards neutrophils in severe patients, consistent with previous findings (*31*). Similar patterns are observed in the Liao dataset where T cell to T cell interaction was higher in moderate patients, while interactions involving neutrophils and monocytes increased in severe patients (Supplementary Fig. 2C).

The enrichment of monocyte dominated interactions in severe patients is also observed in all three PBMC datasets (Supplementary Fig. 5D-F). Interestingly, two of the PBMC datasets (Zhang and Arunachalam) illustrate that severe patients have a decrease in CD8 memory T cell interactions. The lack of T cell interactions in severe patients suggests potential T cell depletion, a finding supported by previous studies that lymphopenia is associated with severe disease (*32*–*34*). Together, these data provide validation that our workflow can confirm known mechanisms and highlight new biology for further investigation.

### Monocyte/macrophage and neutrophil interaction in severe patients are dominated by CXCL, IL1 and other inflammation pathways

Focusing on individual pathways, Fig. 3A illustrates that all pathways can be broadly grouped into six large clusters. In particular, two of these pathway-clusters (pathway-cluster 2 marked by orange and pathway-cluster 4 marked by pink) are dominated by inflammatory pathways and these have significantly higher interaction between monocytes and neutrophils in severe patients compared to moderate (Fig. 3A-B). This is consistent with findings that in the productive immune response to SARS-CoV-2 infection, alveolar macrophages recognize and phagocytize apoptotic cells; however, under a dysfunctional immune response, excessive activation and accumulation of monocytes/macrophages and neutrophils leads to the overproduction of inflammatory cytokines which then damages the lung and other organs (*35, 36*).

**Fig. 3:**
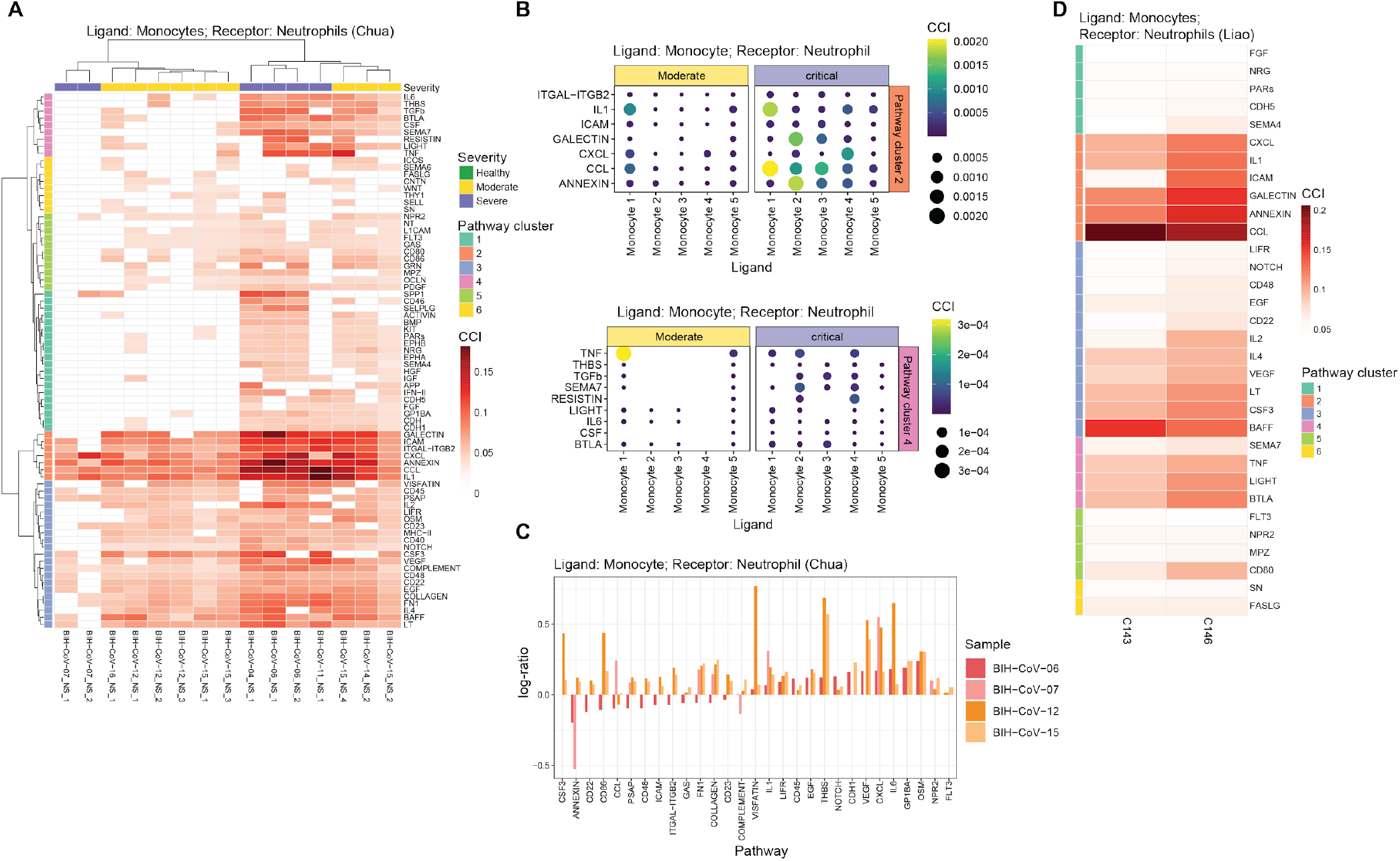
A. Heatmap of the pathway-specific cell-cell interaction (pCCI) contribution in monocytes as ligands and neutrophils as receptors in the Chua dataset, where the rows indicate the signaling pathways and columns indicate the samples. The signaling pathways are clustered into 6 groups. B. Dot plot indicating the cell-cell interaction contribution (pathway-cluster cell-cell interaction) in monocyte subgroups as ligands and neutrophils as receptors of the pathway-cluster 2 (upper panel) and pathway-cluster 4 (lower panel) as defined in (A). The columns indicate the 5 cellular subtypes of monocytes as ligands and the rows indicate the signaling pathways. A larger dot represents a higher level of cell-cell interaction. C. Bar plot indicating the log-ratio of cell-cell interaction contributions between two time points (y-axis) for longitudinal samples of 4 patients (2 moderate: BIH-CoV-12, BIH-CoV-15; 2 severe: BIH-CoV-06, BIH-CoV-07) in monocytes as ligands and neutrophils as receptors. The x-axis represents the signaling pathways. D. Heatmap of the cell-cell interaction contribution in monocytes as ligands and neutrophils as receptors in the Liao dataset, where the rows indicate the signaling pathways and columns indicate the samples. The signaling pathways are highlighted by the 6 signaling pathway clusters from (A).

To further delineate differences between moderate and severe patients observed in Fig. 2D (shown by thick red edges between monocytes and neutrophils), we investigated which subpopulations of monocytes actively interact with neutrophils (Supplementary Fig. 6A-B). The two inflammatory pathway-clusters mentioned above show that different cellular subtypes of monocytes in severe patients have significantly higher interaction scores in different pathways (Fig. 3B). More specifically, we found that in severe patients, cellular subtype “monocyte 1” interacts with neutrophils through IL1, and CCL pathways, whereas interactions in moderate patients are dominated instead by TNF. Supplementary Fig. 6C-D shows that “monocyte 1” is marked by genes IL1B, IL1RN, IL8, TNFRSF1B and CCL4 and characterized by gene ontology terms “regulation of inflammatory response” as well as “regulation of apoptotic signaling pathways”. The cellular subtype “monocyte 2” (marked by highly expressed IFI27), interacts primarily with neutrophils through pathways ANNEXIN and GALECTIN, which could suggest a role for this cluster in phagocytizing dying neutrophils. The cellular subtype “monocyte 3” expressing IFIT2, IFIT3, CCL8, CXCL10, and CXCL11 shows strong signatures of type 1 interferon cell-cell signaling (Fig. 3B and Supplementary Fig. 6C), suggesting equal support for antiviral immunity in moderate and severe patients. Alternatively, proinflammatory signaling via CXCL interactions is mainly through cellular subtype “monocyte 4”, which highly expresses CCL2, CXCL1, CXCL2 and CXCL5 (Fig. 3B and Supplementary Fig. 6C).

Similar patterns are observed in the monocyte-neutrophil interaction in BALF (*24*) tissues where patient samples with neutrophils have higher interaction signaling from monocytes through pathways CXCL, IL1, GALECTIN, ANNEXIN, and CCL (Fig. 3D) demonstrating the consistency of our cell-cell interaction results across nasopharyngeal and bronchoalveolar lavage fluid samples. The impact of CXCL and IL1 are also found among the four sets of longitudinal samples in the Chua dataset under different disease progression, suggesting an increase in interactions of signaling pathways CXCL, IL1 over time (Fig. 3C). Interestingly, ANNEXIN downregulates across time in severe patients, but increases in moderate patients (Fig. 3C).

### Interaction from goblet cells to immune cells are heterogeneous in moderate patients and severe patients

We observe heterogeneous interaction patterns from goblet cells to immune cells across patients and pathways (Fig. 4A). Goblet cells are found to express high levels of genes associated with innate and antiviral immune functions indicating that the nasal epithelial cells interacting with immune cells may play an important role in reducing early viral load and this is also consistent with recent literature (*37*). We observe one subgroup of severe patients (n = 3; including one deceased patient) showing clear differences in cell-cell interaction within the pathway-cluster 1 compared to moderate patients. In particular, they show a lack of interaction in the collection of pathways which includes immune signaling and costimulation pathways such as CD40, CD80, CD23 and CD86 inflammatory pathways IL6, IFN-II, and Th2 cytokines IL-4/IL10 (Fig. 4A). Another subgroup of severe patients (n = 6) show clusters with a small subgroup of moderate patients that has low cell-cell interaction for antigen presentation (MHC-II), signaling pathways PTN and NPR2, and this subgroup is also lacking the Th2 cytokine IL-4 and the B cell activating factor BAFF (Fig. 4A). Together, these results point to cohort heterogeneity within severe patients implicating immune costimulation or T cell polarizing pathways may contribute to disease severity with context.

**Fig. 4:**
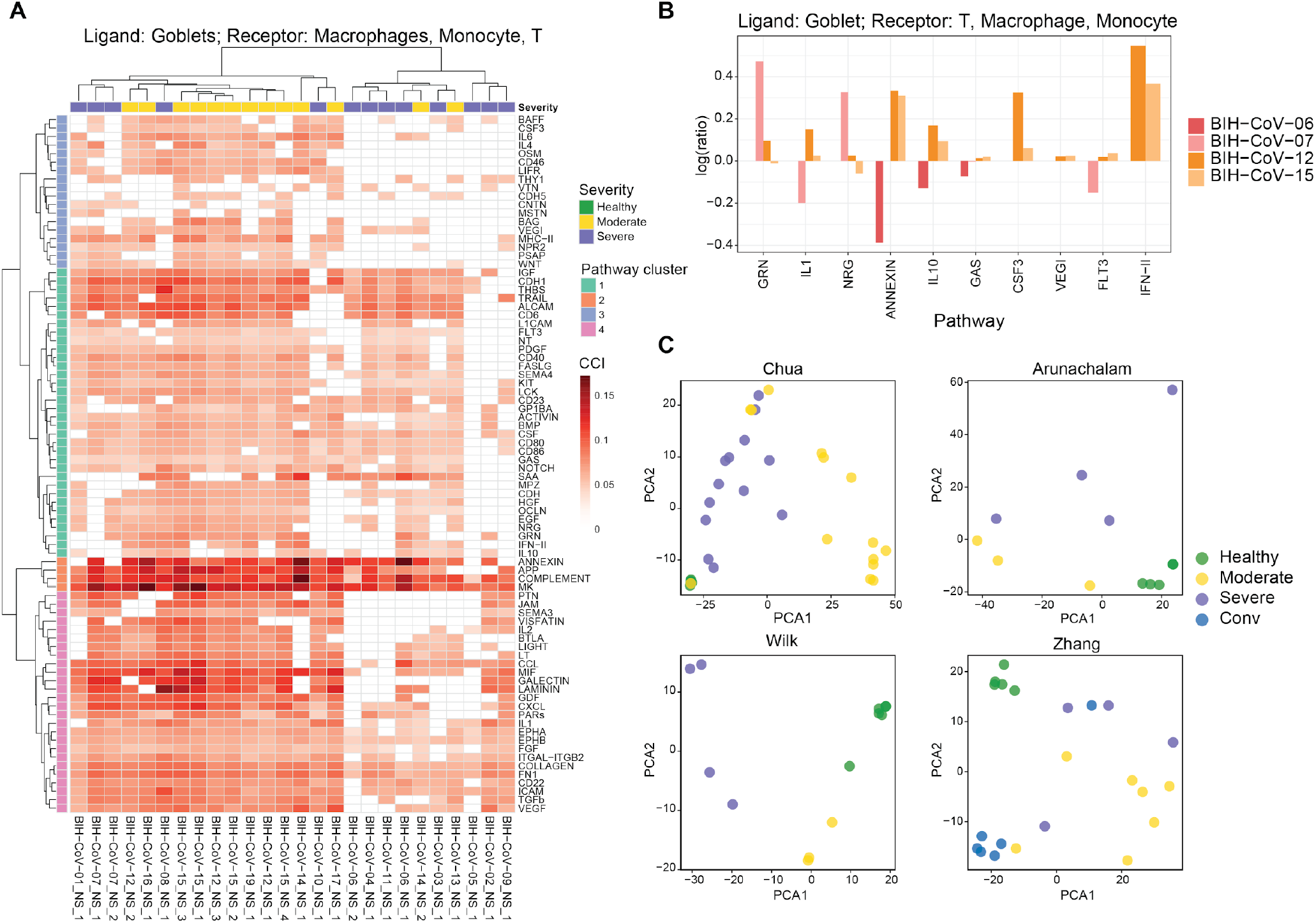
A. Heatmap of the pathway-specific cell-cell interaction contribution in goblets as ligands and immune cells (macrophages, monocytes and T cells) as receptors in the Chua dataset, where the rows indicate the signaling pathways and columns indicate the samples. The signaling pathways are clustered into 6 groups. B. Bar plot indicates the log-ratio of cell-cell interaction contributions between two time points (y-axis) for longitude samples of 4 patients (2 moderate: BIH-CoV-12, BIH-CoV-15; 2 severe: BIH-CoV-06, BIH-CoV-07) in goblets as ligands and immune cells (macrophages, monocytes and T cells) as receptors. The x-axis represents the signaling pathways. C. PCA for samples using the selected pathway-specific cell-cell interaction features, colored by disease severity: the Chua dataset (top left panel), the Wilk dataset (bottom left panel), the Arunachalam dataset (top right panel) and the Zhang dataset (bottom right panel) with the corresponding LOOCV accuracy rate for four datasets presented in Supplementary Table 1.

Focusing on a specific cellular subtype of epithelial cells (goblet 5), we observe a number of increased activities in moderate patients in the ANNEXIN pathway. This is most evident between cellular subtypes “goblet 5” and “monocyte 5”. This epithelial to immune cell interaction within the ANNEXIN pathway also shows an increase in patients under moderate conditions (Fig. 4B). Annexin plays a role in phagocytic uptake of dying cells, can drive neutrophil detachment and apoptosis, and plays a predominant role in immune resolution (*38*). Remarkably, glucocorticoids, which are effective at treating COVID-19 patients, act via upregulating Annexin I (*39*), suggesting that natural moderate symptoms for COVID-19 may be linked to effective endogenous immune management, or that patients that respond to glucocorticoid drugs elevate Annexin cell communication pathways that then limit further inflammation, and this response is detectable via single-cell analysis. We have developed and provide an interactive resource (http://shiny.maths.usyd.edu.au/CovidCellInteraction/) to enable further investigation of cell-cell interaction at different resolutions from aggregate interaction between two major cell types to visualize expression values for specific ligand-receptor pairs.

### Cell-cell interaction patterns have the potential to discriminate between moderate and severe patients

Finally, we found that information from cell-cell interactions provides a discriminating signal for patients with different disease progression. Fig. 4C shows the principal components of the cell-cell interaction matrix for the Chua dataset with samples from healthy controls, moderate and severe patients highlighted. Linear discriminant analysis (LDA) shows that based on accuracy, ligand-receptor features selected based on interaction from epithelial (ciliated and goblet) cells to immune cells (LOOCV = 0.8) has a higher discriminating power than using cell type proportion (LOOCV = 0.4) or ligand and receptor gene expression alone (LOOCV = 0.6). Examples of top selected pathways are THBS, BMP and EGF from pathway-cluster 1, and MHC-II and COMPLEMENT from pathway-cluster 2 (pathway-cluster defined in Fig. 4A). This result is consistent regardless of the statistical machine learning methods employed (Supplementary Table 1). The accuracy rate of leave-one-out cross-validation (LOOCV) based on the first three PCs using k nearest neighbor classification (k = 3) is 84.4%, highlighting the ability of cell-cell interaction features to predict the degree of severity of patients. By repeating our workflow on the three PBMC datasets, we further demonstrate that using CCI features can achieve higher LOOCV accuracy rate than using cell type composition as features. Despite the limited samples, repeating our workflow on a smaller dataset within BALF tissues in the Liao dataset demonstrates similar findings that cell-cell communication patterns between goblet cells to immune cells has potential discriminating power.

## Discussion

A better understanding of virus and host cell interaction at the cellular level is an important component in understanding infectious disease progression and is critical for developing a treatment for the disease. In this paper, we provide a comprehensive workflow to integrate and examine multiple COVID-19 single-cell RNA-seq datasets to identify differential cell-cell interaction (CCI) pathways with respect to disease. Our initial results in upper airway tissues show strong intra-epithelial communication in the healthy lung, whereas the immune system then dominates communication pathways during COVID-19. We then discover that despite a higher cell-cell interaction (tCCI score) in severe patients compared to moderate patients between immune and neutrophil cells, the CCI scores between epithelial and immune cells are heterogeneous among severe patients, with a subpopulation illustrating lower CCI score when compared to moderate patients. Furthermore, features extracted from cell-cell interactions are potential signatures for discriminating between moderate and severe patients.

In most multi-omics profiling in patients with COVID-19, strong acute inflammatory responses are commonly found in most of the cell types as expected. Since the airway epithelium is the primary site of infection for SARS-CoV-2 causing disease, investigating how epithelial cells interact with immune cells differentially leads to a better understanding of the initial host reaction to viral infection. Therefore, examining cell-cell communication offers an analytical approach to characterize specific cell type interaction and identify potential immune response drivers that results in different degrees of disease severity.

The importance of using a workflow that accounts for cohort heterogeneity in examining severe and moderate patients is clearly illustrated when we examine the interaction pattern between nasal epithelium as a ligand and various receptors in immune cells. This is a different approach to the one taken by Chua and colleagues (*25*), where they measure the overall/aggregate interaction between epithelial and immune cells and found a higher overall/aggregate interaction between epithelial and immune cells in severe patients. Here, when we examine the cell-cell interaction relationships at the individual sample level, we observe clear cohort heterogeneity among severe patients, and we are able to discover a subgroup of the moderate patients with higher interaction between epithelial and immune cells.

In this study, we focus on the cell communication within COVID-19 patients via ligand-receptor signaling. Several methods have been developed recently to infer such cell-cell interaction from scRNA-seq data, such as CellPhoneDB, SingleCellSignalR, NicheNet and CellChat (*20*–*22, 40*). Most of these methods aim to identify the significant ligand and receptor gene pair between two cell populations with the most recent method CellChat (*22*) that accounts for additional signaling factors. In addition, CellChat systematically categorizes the ligand-receptor pairs based on their signaling pathways, providing a comprehensive interpretation of cell-cell communication from single-cell RNA-seq. There are also other types of cell communication like physical cell interaction that can be further investigated. Technology to sequence physically interacting cells like PIC-seq has been used to investigate epithelial–immune interaction and infectious disease in mice (*41*). Application of such technology in COVID-19 research will potentially allow characterization of differential physical intercellular interaction at high resolution.

Our analysis suggests the heterogeneity of cell-cell interaction patterns within patients, even if they have similar degrees of symptoms. One key variability is the sampling time since the onset of symptoms, as this may not fully capture the true underlying disease progression within each individual. Other potential factors that lead to the variability include age, gender, comorbidities and viral load.

Currently, with the limited number of samples from patients with similar clinical characteristics, accounting for these uncertainties in modelling is challenging. Towards the future, as more large single-cell profiling resources in COVID-19 become publicly available, integrative analysis and meta-analysis of these studies by incorporating patient diversity to our workflow will provide a more comprehensive characterization of cell-cell interaction patterns in COVID-19 patients. Nevertheless, using the current databases our workflow supports that cell-cell interactions provide more meaningful predictions of disease progression (Figure 4C).

In summary, our novel workflow enables integrative analysis of five different COVID-19 scRNA-seq data sets with a total of 415,856 cells and 85 samples. This generalizable workflow was built on the latest single-cell analytical methods and enables the identification of differential cell-cell interaction across disease progression. We discover clear cohort heterogeneity among the severe patients in the interaction between epithelial and immune cells, with signatures that can be linked with patient outcome. Together we provide a validated workflow for integration and analysis of diverse single-cell sequencing data to pinpoint communication networks that control disease outcome.

## Materials and Methods

### Data and preprocessing

[A]Chua dataset - The raw count matrix and metadata containing patient information are downloaded from FigShare: https://doi.org/10.6084/m9.figshare.12436517 (*25*). This data includes 19 patients with critical or moderate disease as well as 5 healthy controls.

[B] Liao dataset - The raw count matrices of single-cell RNA-seq data from bronchoalveolar lavage fluid was downloaded from the National Center for Biotechnology Information (NCBI) Gene Expression Omnibus (GEO) under the accession number GSE145926. This data has 3 healthy controls, 3 moderate patients and 6 severe patients (*24*).

[C] Wilk dataset - The raw count matrices of single-cell RNA-seq data from PBMC with metadata were downloaded from the COVID-19 Cell Atlas: https://www.covid19cellatlas.org/#wilk20 (*28*). This data contains 6 healthy controls, 3 moderate patients and 4 severe patients.

[D] Arunachalam dataset - The raw count matrices of single-cell RNA-seq data from PBMC and the clinical information were downloaded from GEO under accession number GSE155673. This data has 5 healthy controls, 3 moderate patients and 4 severe patients (*29*). The cells with more than 20% mitochondrial proportion and UMI count greater than 50,000 are removed from the downstream analysis.

[E] Zhang dataset - The raw sequence files of single-cell RNA-seq data from PBMC are downloaded from the Genome Sequence Archive of the Beijing Institute of Genomics (BIG) Data Center, BIG, Chinese Academy of Science using the accession code HRA000150 (*18*). Cell Ranger (v3.0.2) with human reference version GRCh38 were used to generate the raw count matrices. The data includes 5 healthy controls, 7 moderate patients and 4 severe patients. Only the cells retained from the original study are used.

#### Processing

For each dataset, we performed size factor standardization and log transformation on the raw count expression matrices using the `logNormCount` function in the R package `scater` (version 1.16.2) and generated log transformed gene expression matrices for analysis.

### Computational workflow

#### Step 1 - Cell type annotation

For a given dataset, we perform a cell type identification using the scClassify framework (*19*). Specifically, to identify the cell types from the Chua dataset and the Liao dataset, we performed a modified version of the joint classification from scClassify that incorporates the concept of iterative supervised learning. The initial model is built from four reference datasets including annotated cell information from healthy human lungs (*4, 26, 27*). The final cell type labels were determined by the majority vote from individual classification labels using each single reference. An additional scClassify model based on the assigned cells was then built to predict the cells that are classified as “intermediate” or “unassigned” in the previous step. To identify cell types from the PBMC datasets, we used the Wilk dataset as a reference (*28*) to build the model and use it to predict the cell types for the Zhang dataset and the Arunachalam dataset.

#### Step 2 - Unsupervised clustering for subpopulation identification

We performed unsupervised clustering on each classified cell type to identify the cellular subtypes in the Chua dataset and the Liao dataset. For each cell type, we first calculate the deviance across cells within each sample using the function `devianceFeatureSelection` implemented in the R package `scry` (version 1.0.0). Next, we select features that are among the top 1000 largest deviances in more than 50% of the samples. We then performed negative binomial generalized principal component analysis (GLM-PCA) on the UMI matrix with the selected features (number of components is set to 30) (*42*). A shared nearest neighbor graph is then built based on the GLM-PCA low-dimensional space and used as an input for Louvain clustering to identify subclusters, considering each of them as a refined cellular subtype.

To prevent over clustering, we follow a similar workflow described in clusterExperiment to collapse the identified subclusters (*23*). Hierarchical clustering is first performed on the aggregated average expression of each subcluster to construct a cluster hierarchy, and then from the bottom to top, the clusters of the same branches are merged if less than 10 genes are differentially expressed (log fold change > 1, FDR < 0.01). Note that we identified some cellular subtypes (ionocytes and squamous) that are inconsistently annotated between the original Chua dataset and scClassify (classified as goblet cells). In this instance, based on marker expression, we manually reannotated these two cell types using the original annotation for the downstream analysis.

#### Step 3 - Calculating cell-cell interaction (CCI)

For a given individual sample and a pair of subclusters (i.e. cellular subtypes) obtained in Step 2, we calculate the aggregated ligand-receptor interaction score based on CellChat (*22*) for each signaling pathway. This represents the communication probabilities among all pairs of subclusters across all ligand-receptor pairs. The CellChat algorithm aims to identify the significant ligand-receptor gene pairs between two cell populations while accounting for important signaling factors, including the expression of soluble agonists, antagonists, and stimulatory and inhibitory membrane-bound co-receptors. Finally, the communication probability of a signaling pathway is defined as the sum of the probabilities of its ligand-receptor pairs.

The implementation is available as R code stored at the GitHub, https://github.com/SydneyBioX/COVID_CCI_analysis and as a web shiny application at http://shiny.maths.usyd.edu.au/CovidCellInteraction/.

### Statistical formulation

The <u>output</u> of the cell-cell interaction analysis can be considered as a three-dimensional array representing the cell-cell interaction (CCI) score. Let *x*_*cpk*_ denote the cell-cell interaction (CCI) score generated from the computational workflow for a pair of cellular subtypes *c*, where *c* ∈ *C* (defined below as a set consisting of all pairs of cellular subtypes), pathway *p* with *p* = 1, …, *P*, and individual sample *k* with *k* = 1, …, *K*. In general, an individual sample *k* represents the sample from one individual collected at a specific time point.

For *N* major cell types, we denote them by the sets *M*_1_, *M*_2_, …, *M*_*N*_ and within a given major cell type *M*_*i*_ consisting of *n*_*i*_ cellular subtypes 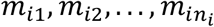 we can write *M*_*i*_ = {*m*_*iu*_|*u* = 1, … *m, n*_*i*_}. We can represent a pair of cellular subtypes as *c* = (*m*_*iu*_, *m*_*jv*_), where *i, j* = 1, …, *N*; *u* = 1,. ., *n*_*i*_ and *v* = 1, …, *n*_*j*_. Here, we consider *m*_*iu*)_ as the sender cellular subtype within major cell type *M*_*iu*_ and *m*_*jv*_ as the receiver cellular subtype from major cell type *M*_*j*_. The collection of all pairs of cellular subtypes is written as *C* = {(*m*_*iu*,_*m*_*jv*_)|*i, j* = 1, …, *N*; *u* = 1, …, *n*_*i*_; *v* = 1, …, *n*_*j*_=. We further denote 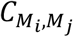 as a subset of *C* containing only pairs of cellular subtypes from major cell type *M* to *M*_*j*_ which is represented as 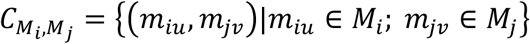.

For a given sample *k*, the following measures of interest are explored:

- Subtype cell-cell interaction (sCCI) between a pair of cellular subtypes *c* for an individual sample *k* is calculated as sCCI (*c, k*)= ∑_*p*_ *x*_*cpk*_. This measure totals the cell-cell interaction score across all pathways. Calculating this score for each pair of cellular subtypes and each individual sample is the same as totaling the array *X* across the pathways resulting in a |*C*| × *K* two-dimensional matrix.
- Pathway specific cell-cell interaction (pCCI) from the major cell type *M*_*i*_ to the major cell type *M*_*j*_ for a pathway *p* and an individual sample *k* is 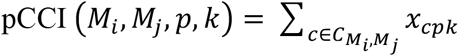 where 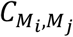 is defined as above. This is a measure that sums the cell-cell interaction scores across all cellular subtypes between any two major cell types. For each pair of (*M*_*i*_, *M*_*j*_), calculating this statistic for each pathway *p* and individual sample *k* results in a *P* × *K* matrix (see Figure 3A).
- Total CCI (tCCI) from major cell type *M*_*i*_ to major cell type *M*_*j*_ for an individual sample *k* is defined as 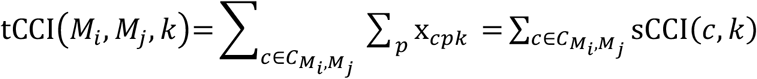, where 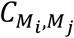is defined as above. This is a measure that sums the cell-cell interaction scores across all cellular subtypes between two major cell types and across all pathways. For each individual sample *k*, calculating the tCCI statistic for each pair of (*M*_*i*_, *M*_*j*_) will result in a *N* × *N* matrix that can be visualized as a heatmap or network graph.
- Suppose 𝒫 represents a set of pathways belonging to the same cluster termed as a pathway-cluster (see Clustering in Methods). The pathway-cluster cell-cell interaction for an individual sample *k* between a pair of cellular subtypes is defined as 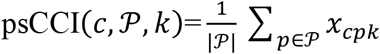.

### Association analysis for CCI

We calculate a group specific cell-cell interaction (CCI_group_) between two cellular subtypes where groups represent any treatment of interest. Here it refers to control and disease progression such as moderate and severe patients. Let 𝒦_group_ denote a set of individual samples under the same condition of interest, where |𝒦_group_| indicates the size of the set. For example, the total number of samples having moderate response to COVID-19 in the dataset (see Fig. 2 A-C). The CCI_group_ from the major cell types *M*_*i*_ to the major cell types *M*_*j*_ can be calculated by 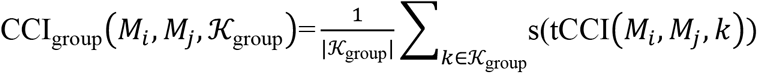, where 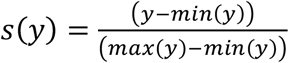 scaling function to scale between individual samples. In practice, the **differential CCI** from *M*_*i*_ to *M*_*j*_ between moderate (CCI_moderate_) and severe (CCI_severe_) patients can be calculated by CCI_severe_ − CCI_moderate_ measuring the differential patterns of the cell-cell interaction across different disease severity (see Figure 2D).

The pathway-cluster cell-cell interaction (used in Figure 3B) for a group of individuals ℒ between a pair of cellular subtypes is simply the sums of psCCI across individual with a group ℒ and can be written as 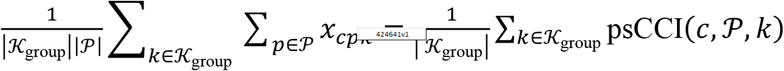. For a pair of cellular subtypes *c*, calculating this statistic results in a | 𝒫 | × |𝒦_group_| matrix.

### Statistical analysis of longitudinal data

Suppose we have multiple samples collected from the same individual at different time points, say *k*_early_ and *k*_late_ then the cell-cell interaction across disease progression is the log-ratio of cell-cell interaction (illustrated in Fig. 3C) between these two time points for a given pair of cell types (sender cell type *M*_*i*_, receiver cell types *M*_*j*_ within a pathway *p* is log (pCCI(*M*_*i*,_ *M*_*j*_, *k*_late_)/pCCI(*M*_*i*,_ *M*_*j*_, *k*_early_)).

### Clustering

We group various pathways based on the similarity of intercellular communication patterns using hierarchical clustering with Euclidean distance and ward.D2 agglomerative method implemented in the function `hclust` in R.

### PMBC data integration

We integrate the three PBMC datasets using a modified version of scMerge (*30*). Here, cell types annotated by scClassify are used as an input to scMerge to construct pseudo-bulk expression profiles. The resulting profiles are used to identify mutual nearest subgroups as pseudo-replicates and to estimate parameters of the scMerge model.

### Machine learning for discrimination

To select the cell-cell interaction features that discriminate across samples under different conditions, we performed a Kruskal-Wallis rank sum test on pathway-specific cell-cell interaction (pCCI) to select the pathways that are significantly different across samples from healthy controls, moderate patients and severe patients. Feature selection is based on pCCI features with an adjusted p-value less than 0.1 for the Chua dataset, less than 0.2 for the Wilk and Zhang datasets and less than 0.4 for the Arunachalam dataset, we termed these selected features as “Top CCI”. For the Chua dataset, we also selected the top pCCI from the cell-cell interaction between the two major epithelial cell types (Goblet and Ciliated) and the immune cell types (B cells, dendritic cells, macrophages, monocytes and T cells), termed as “Epi-Immune CCI”. We further considered cell type proportion as another type of feature.

The classification model to predict the samples’ condition is built with linear discriminant analysis (LDA) and random forest (RF) on the selected features (Top CCI, Epi-Immune CCI, and cell type proportion) as well as k nearest neighbor classification (with k = 1, 3) using the first 3 principal components of the pCCI matrix. The classification performance was determined by leave-one-out cross-validation.

### Gene Ontology analysis

Differential gene expressions were identified using moderated t-statistics implemented in the R package `limma` (version 3.44.3). The gene set over-representation analysis for the significant DE genes (top 100 genes selected) with biological process (BP) gene ontology is measured using the “`enrichGO`” function in the R package `clusterProfiler` (version 3.16.0) (*43*). Significant GO term is defined by q-value < 0.1.

### Interactive graphics implementation

To facilitate the interpretation of the complex data set, we have created an online interactive tool which allows researchers to explore different parts of the data. The first tab of the tool contains four columns. The first column allows the user to select two groups (or individual samples) to compare and it displays the associated cell-cell interaction network. The second column shows the difference between the two selected groups (or samples) in a heatmap and network form. Selecting a cell type pair from the heatmap dissects the interaction into individual pathways and sub-cell types, displayed in the third column. Selecting a pathway on this heatmap further dissects the activity into individual ligand-receptor pairs, displayed in the fourth column. The second tab of the tool allows the user to select a gene and its mean expression is shown for each cell type and sample. The user can also select a ligand cell type and a receptor cell type and the activity of all pathways between these cell types and involving the selected gene are shown.

## Acknowledgments

The authors thank all their colleagues, particularly at The University of Sydney, Sydney Precision Bioinformatics Alliance and Charles Perkins Centre for their support and intellectual engagement.

## Funding

The following sources of funding for each author, and for the manuscript preparation, are gratefully acknowledged: Australian Research Council Discovery Project grant (DP170100654) to JYHY. Research Training Program Tuition Fee Offset and Stipend Scholarship and Chen Family Research Scholarship to YL. The funding source had no role in the study design; in the collection, analysis, and interpretation of data, in the writing of the manuscript, and in the decision to submit the manuscript for publication.

## Author contributions

JYHY and YL conceived the study and developed the workflow. YL processed and curated all the scRNA-seq data with input from JY. YL performed all data analysis with input from all authors. LL, CM, DL & GN jointly examined and analyzed the biological findings. AT implemented the R shiny app with input from all authors. All authors wrote the manuscript and approved the final version of the manuscript.

## Competing interests

The authors declare no competing interests.

## Data and materials availability

All data needed to evaluate the conclusions in the paper are present in the paper and/or the Supplementary Materials.

## Supplementary Materials

**Supplementary Fig. 1.**
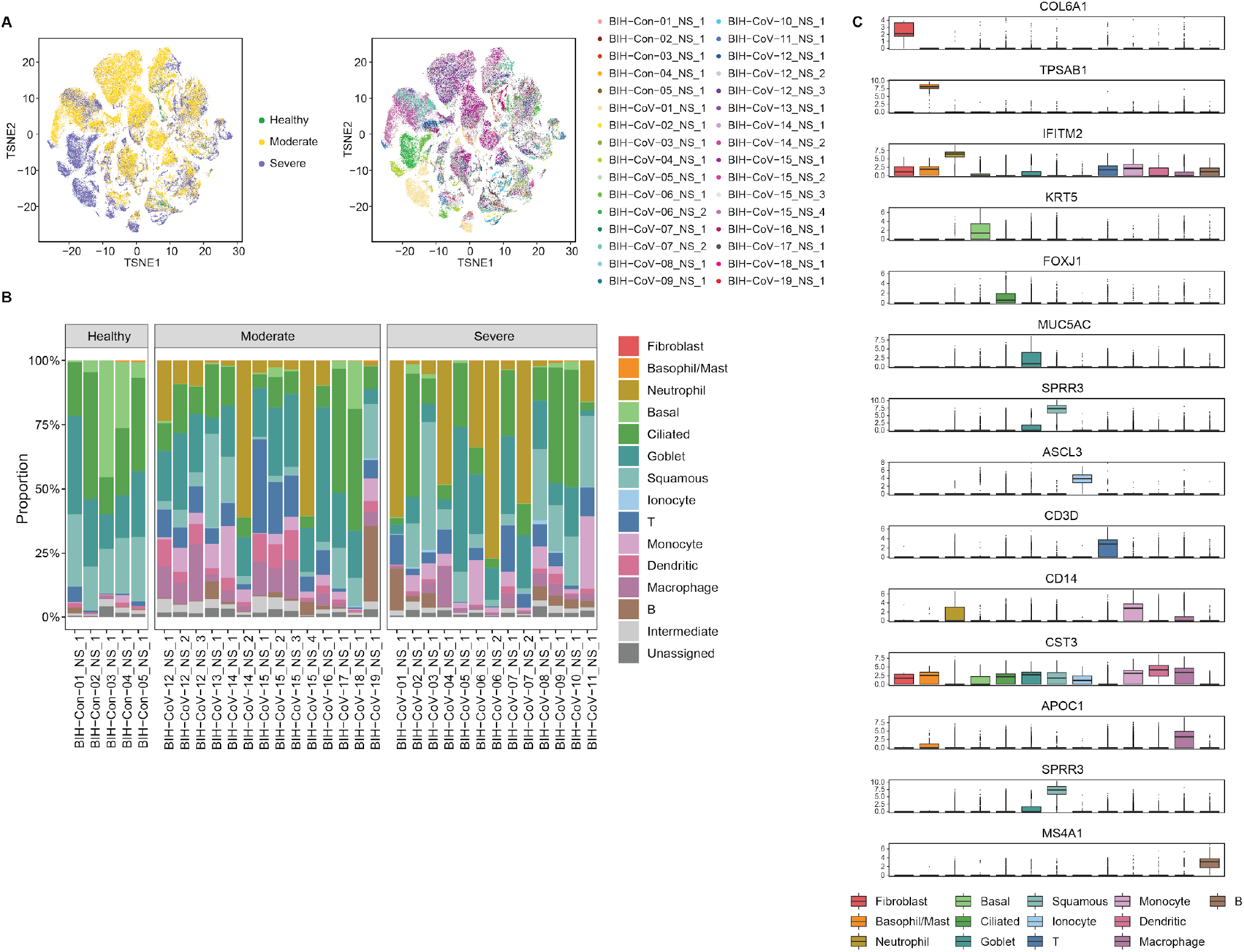
A. tSNE plots with the Chua dataset, colored by the disease condition (left panel), and individual sample (right panel). B. Cell type composition of each individual sample in the Chua dataset. C. Boxplots of marker expression for each reannotated cell type.

**Supplementary Fig. 2.**
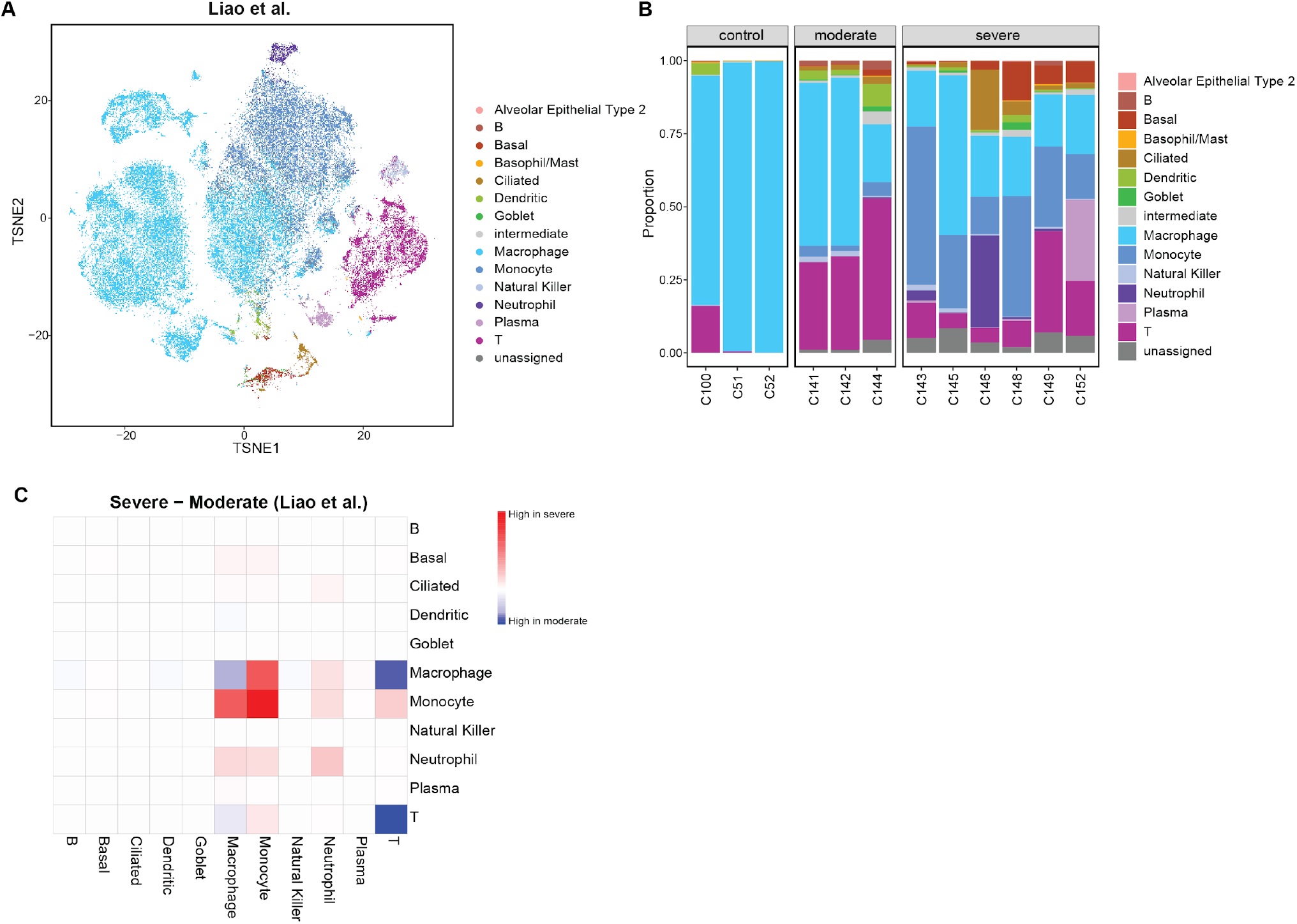
A. tSNE plot of scRNA-seq data from BALF (the Liao dataset), colored by the reannotation from scClassify. B. Cell type composition of each sample in the Liao dataset. C. Heatmap indicating the difference of group specific cell-cell interaction between different cell types in severe patients and moderate patients in the Liao dataset. Red color indicates a higher interaction in severe patients and blue color indicates a higher interaction in moderate patients. Rows indicate the sender cell types and columns indicate the receiver cell types.

**Supplementary Fig. 3.**
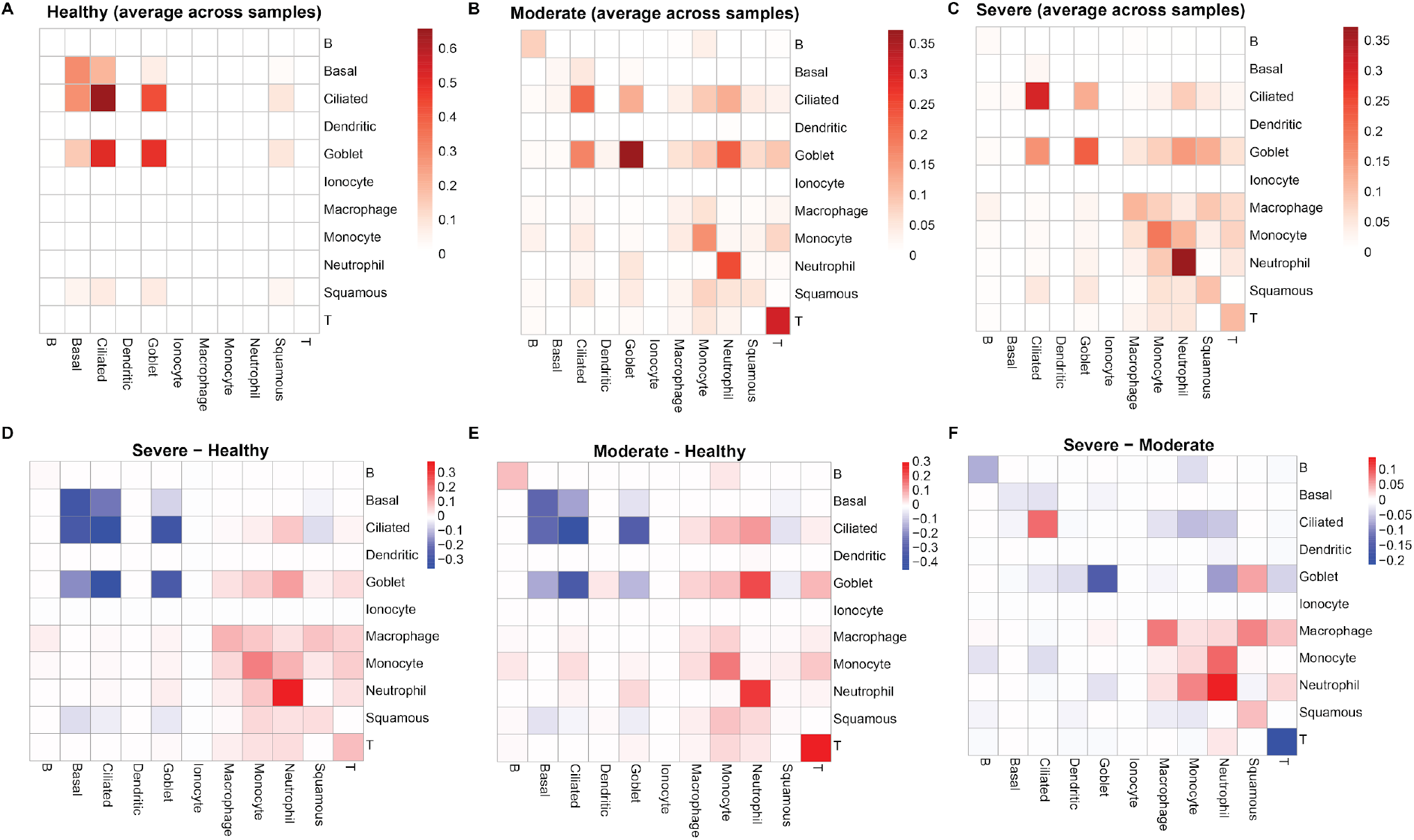
(A-C) Heatmaps indicating the group specific cell-cell interaction between different cell types in (A) healthy controls (B) moderate patients (C) severe patients for the Chua dataset. Rows indicate the sender cell types and columns indicate the receiver cell types. (D-F) Heatmaps indicate the difference in group specific cell-cell interaction between different cell types in (D) severe patients and healthy controls (E) moderate patients and healthy controls (F) severe patients and moderate patients for the Chua dataset. Red color indicates a higher interaction in severe patients and blue color indicates a higher interaction in moderate patients. Rows indicate the sender cell types and columns indicate the receiver cell types.

**Supplementary Fig. 4.**
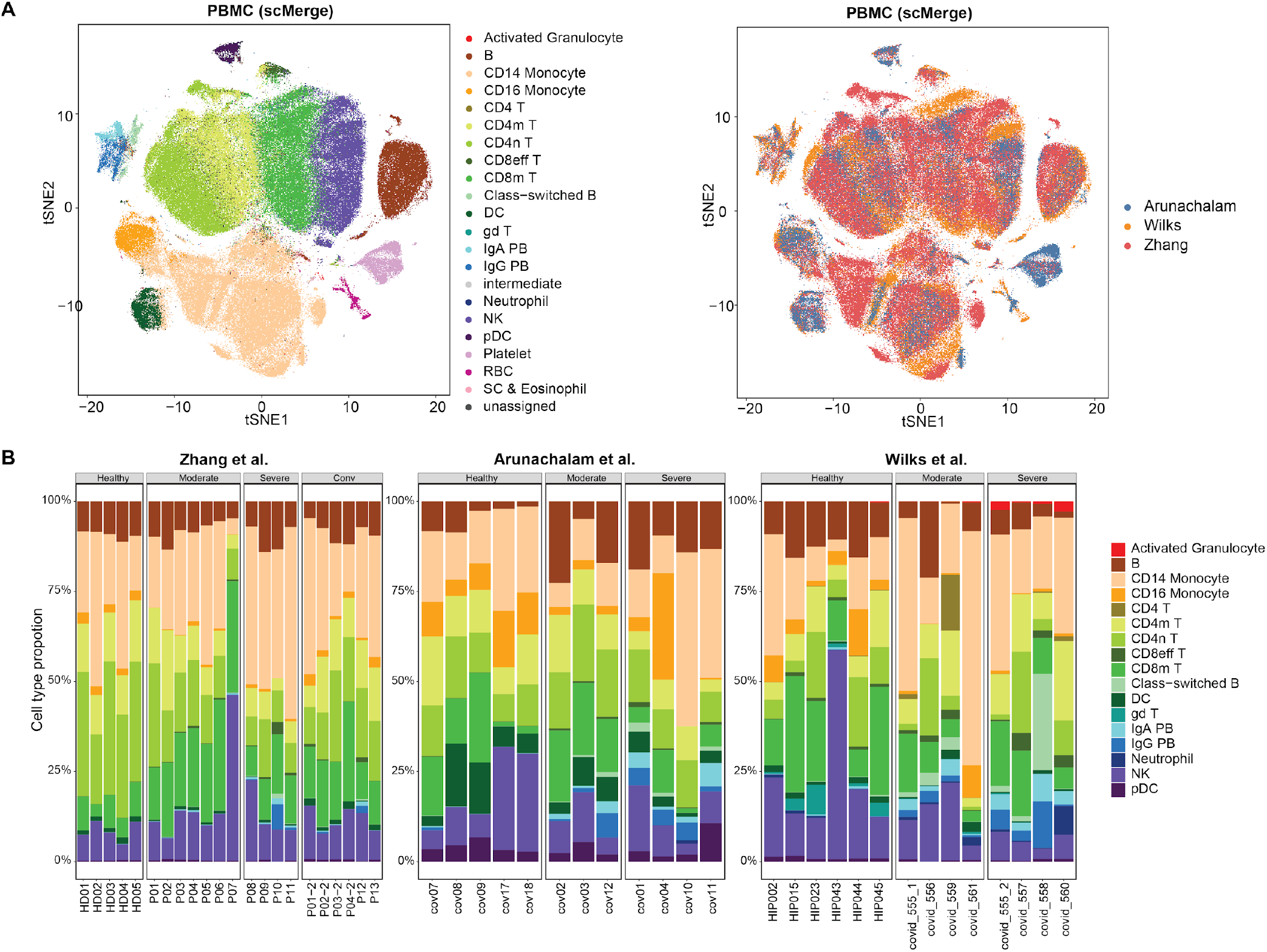
A. tSNE plots of the integrated matrix generated from scMerge for the three PBMC datasets, colored by the reannotation of cellular subtypes from scClassify (left panel), and colored by datasets (right panel). B. Cell type composition of each individual sample in the three PBMC datasets.

**Supplementary Fig. 5.**
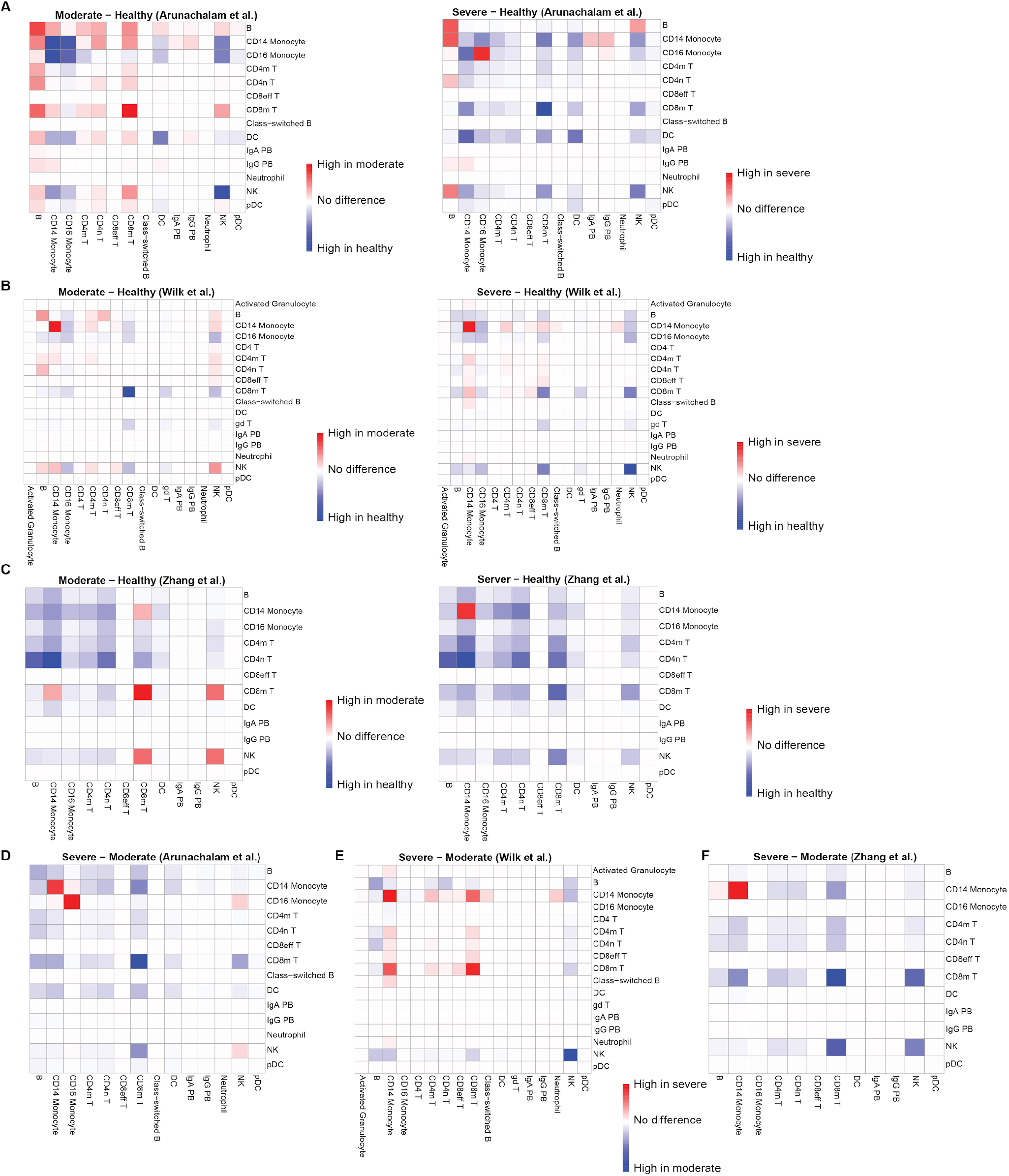
(A-C) Heatmaps indicate the difference of group specific cell-cell interaction between different cell types in moderate patients and healthy controls (left) and severe patients and healthy controls for (A) the Arunachalam dataset (B) the Wilk dataset, and (C) the Zhang dataset. Rows indicate the sender cell types and columns indicate the receiver cell types. (D-F) Heatmaps indicate the difference in cell-cell interaction between different cell types in severe patients and moderate patients for (D) the Arunachalam dataset, (E) the Wilk dataset, and (F) the Zhang dataset. Red color indicates a higher interaction in severe patients and blue color indicates a higher interaction in moderate patients. Rows indicate the sender cell types and columns indicate the receiver cell types.

**Supplementary Fig. 6.**
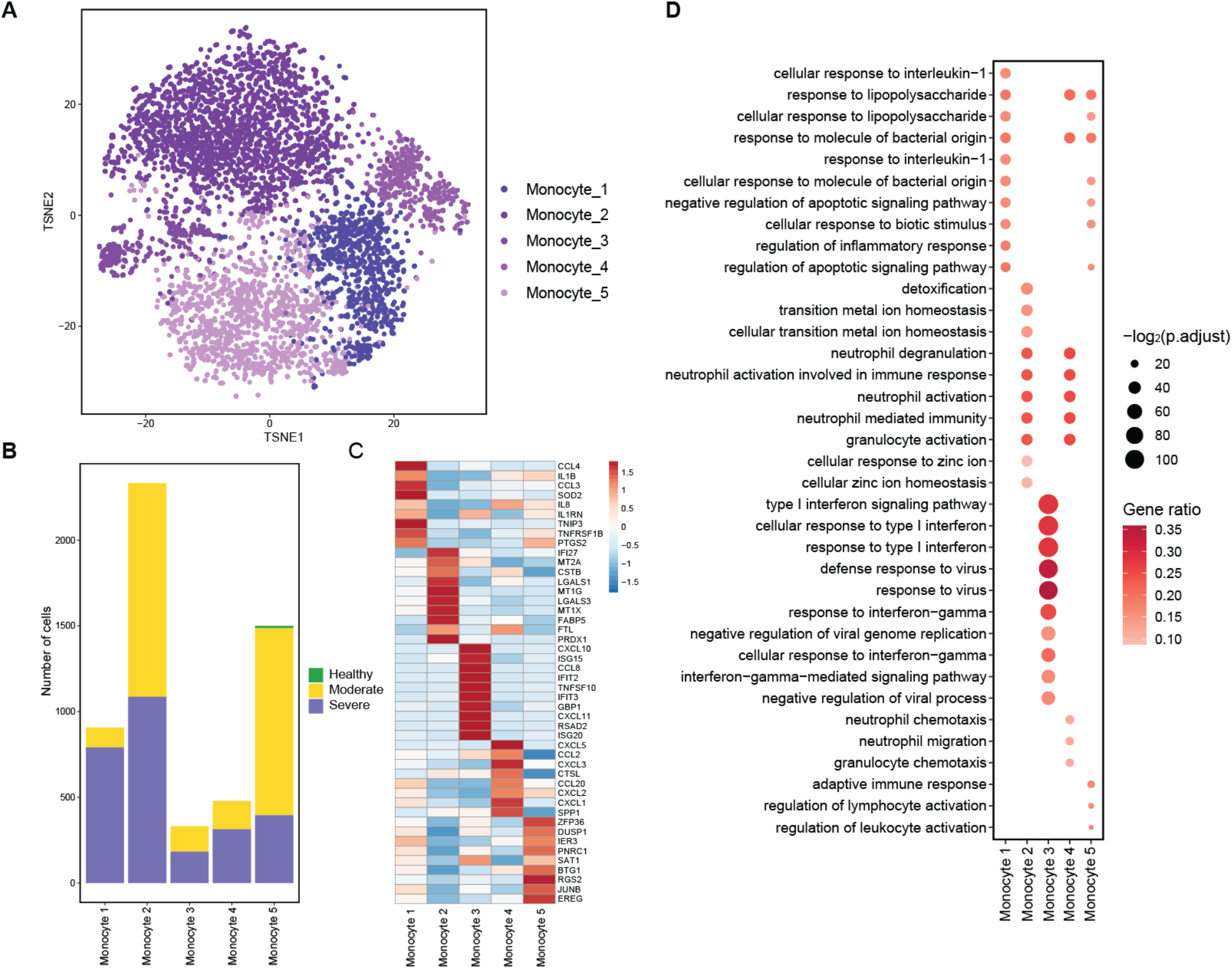
A. tSNE plot of monocytes in the Chua dataset, colored by the five cellular subtypes of monocytes. B. Stacked bar plots representing the number of cells for healthy, moderate and severe groups. The x-axis represents the five cellular subtypes of monocytes for the Chua dataset. C. Heatmap indicates the scaled average marker expression of the five cellular subtypes of monocytes. D. Gene ontology analysis for the cellular subtypes of monocytes.

**Supplementary Table 1.** LOOCV accuracy rate for four datasets using four classification methods: KNN (K = 1), KNN (K = 3), linear discriminant analysis (LDA), and random forest (RF). The row “Top CCI” refers to classification results based on features selected by Kruskal-Wallis rank sum test on pathway-specific cell-cell interaction (pCCI) (See Material and Methods section for more details). The row “Epi-Immune CCI” refers to classification results based on features selected from the cell-cell interaction between the two major epithelial cell types (Goblet and Ciliated) and the immune cell types (B cells, dendritic cells, macrophages, monocytes and T cells). The row “cell type proportion” refers to classification results based on the cell type proportion. The highlighted cells indicated the best performing signature(s) for each of the classification methods.

Chua
|  | KNN (K = 1) | KNN (K = 3) | LDA | RF |
| --- | --- | --- | --- | --- |
| Top CCI | 0.75 | 0.72 | 0.62 | 0.66 |
| Epi-Immune CCI | 0.81 | 0.72 | 0.78 | 0.72 |
| Cell type proportion | 0.47 | 0.5 | 0.44 | 0.56 |

Arunachalam
|  | KNN (K = 1) | KNN (K = 3) | LDA | RF |
| --- | --- | --- | --- | --- |
| Top CCI | 0.83 | 0.67 | 0.50 | 0.67 |
| Cell type proportion | 0.67 | 0.42 | 0.67 | 0.75 |

Wilk
|  | KNN (K = 1) | KNN (K = 3) | LDA | RF |
| --- | --- | --- | --- | --- |
| Top CCI | 0.92 | 0.92 | 1.00 | 0.77 |
| Cell type proportion | 0.46 | 0.46 | 0.62 | 0.69 |

Zhang
|  | KNN (K = 1) | KNN (K = 3) | LDA | RF |
| --- | --- | --- | --- | --- |
| Top CCI | 0.68 | 0.73 | 0.82 | 0.82 |
| Cell type proportion | 0.36 | 0.27 | 0.64 | 0.68 |

## Notes

https://github.com/SydneyBioX/COVID_CCI_analysis

http://shiny.maths.usyd.edu.au/CovidCellInteraction/.

### Competing Interest Statement

The authors have declared no competing interest.

